# Prediction of Hand Movement Speed and Force from Single-trial EEG with Convolutional Neural Networks

**DOI:** 10.1101/492660

**Authors:** Ramiro Gatti, Yanina Atum, Luciano Schiaffino, Mads Jochumsen, José Biurrun Manresa

**Affiliations:** Laboratory for Rehabilitation Engineering and Neuromuscular and Sensory Research, Faculty of Engineering, National University of Entre Ríos, Oro Verde, Argentina; Institute for Research and Development in Bioengineering and Bioinformatics, CONICET-UNER, Oro Verde, Argentina; Center for Neuroplasticity and Pain, SMI^®^, Aalborg University, Aalborg, Denmark; Center for Sensory-Motor Interaction, SMI^®^, Aalborg University, Aalborg, Denmark

**Keywords:** Convolutional neural networks, hand movement, movement prediction, multi-class classification, single-trial EEG

## Abstract

Building accurate movement decoding models from brain signals is crucial for many biomedical applications. Decoding specific movement features, such as speed and force, may provide additional useful information at the expense of increasing the complexity of the decoding problem. Recent attempts to predict movement speed and force from the electroencephalogram (EEG) achieved classification accuracy levels not better than chance, stressing the demand for more accurate prediction strategies. Thus, the aim of this study was to improve the prediction accuracy of hand movement speed and force from single-trial EEG signals recorded from healthy volunteers. A strategy based on convolutional neural networks (ConvNets) was tested, since it has previously shown good performance in the classification of EEG signals. ConvNets achieved an overall accuracy of 84% in the classification of two different levels of speed and force (4-class classification) from single-trial EEG. These results represent a substantial improvement over previously reported results, suggesting that hand movement speed and force can be accurately predicted from single-trial EEG.

## 1. Introduction

Decoding brain signals to predict movements is useful in many research areas, such as neuromechanics, neuroscience and robotics [1]. Furthermore, it is also relevant in neurological rehabilitation, since it has potential to facilitate the assessment of the central nervous system in patients, promote neural plasticity, improve motor dysfunction and allow the control of assistive devices through brain-computer interfaces (BCI) [2]. In this regard, motor commands generated prior to or during movement execution can be extracted from specific oscillatory patterns in the electroencephalogram (EEG) [3, 4]. Specifically, the component waves of movement-related cortical potentials (MRCPs) immersed in the EEG, such as the readiness potential and contingent negative variation, carry information about anticipatory behaviour, which can be used to predict movements, i.e., to detect and classify a particular movement before it is actually executed during self-paced or cue-based paradigms [5, 6, 7].

The movement decoding process is generally focused on detecting a predetermined final state and lacks attention regarding the quality of the action, resulting in simple, rough commands [8]. Research on fine movements of body structures such as fingers [9], or complex movement control [10] is comparatively scarce. It is straightforward to hypothesise that better commands can be achieved if movement kinematics and kinetics are taken into account in the decoding process [11]. Indeed, the decoding of hand movement velocities [12, 13] and 3D trajectories [14] as well as the prediction of force and speed from a specific movement [15, 10] showed promising results. However, recent attempts to predict speed and force from a hand grasping tasks resulted in a classification accuracy not better than chance level [16, 17, 18], stressing the need for more accurate prediction schemes.

The control strategies generated by the nervous system for goal-directed motor behaviour are extremely complex. Thus, pattern recognition systems used to decode and predict movements require careful engineering and domain expertise to transform raw EEG signals (usually by means of a feature extraction subsystem) into a suitable representation for the classification stage [19]. In this regard, several techniques have been proposed for feature extraction, e.g., common spatial patterns, independent component analysis, and joint time-frequency analysis, and also for classification, e.g., nearest neighbour classifier, linear discriminant analysis, support vector machines (SVMs), and ensemble strategies, among others [20]. An alternative is to use representation learning methods that automatically perform a feature extraction and classification through optimisation algorithms. Deep learning is a paradigmatic example, with multiple levels of representation obtained by combining simple but non-linear modules that transform the input into increasingly more abstract levels [19]. In line with this, a decoding model based on deep learning implemented through convolutional neural networks (ConvNets) recently showed promising results in classification performance using different EEG paradigms [21].

The aim of the present study was to improve the prediction accuracy of hand movement speed and force from single-trial EEG signals recorded from healthy volunteers. Subjects executed an isometric right hand palmar grasp task using two predefined levels of force (20% and 60% of the maximum voluntary contraction, MVC) and speed (a 3-s slow grasp and a 0.5-s fast grasp). EEG data were minimally pre-processed, in order to minimise user bias. A prediction strategy using ConvNets was implemented and contrasted with results obtained using state-of-the-art prediction strategies on the same datasets. Overall classification accuracy, precision, recall and Cohen’s kappa (*κ*) values were quantified in order to evaluate the performance of the proposed prediction strategies.

## 2. Materials and methods

### 2.1. Dataset

A dataset consisting of EEG recordings from sixteen healthy volunteers was employed [16]. Written informed consent was obtained from all subjects prior to participation, and the Declaration of Helsinki was respected. The study was approved by the local ethical committee of Region NordJylland (approval no. N-20100067). EEG was recorded during four isometric right palmar grasp tasks with different execution speeds and force levels (expressed as percentage of MVC), categorised as follows: *Slow20*, 3 s to reach 20% MVC; *Slow60*, 3 s to reach 60% MVC; *Fast20*, 0.5 s to reach 20% MVC and *Fast60*, 0.5 s to reach 60% MVC. Forty externally cued repetitions (trials) were performed for each task. A Neuroscan NuAmp Express amplifier was used to record the EEG (Compumedics Ltd., Victoria, Australia) from the electrode locations shown in Fig. 1, in accordance to the 10/10 system. The corresponding EEG channels were referenced to the right earlobe and grounded at nasion. During the experiment, the impedance of all electrodes was kept below 5 kΩ, continuously sampled at 500 Hz and stored for offline analysis. For additional details of the experimental procedure, please refer to [16].

**Figure 1:**
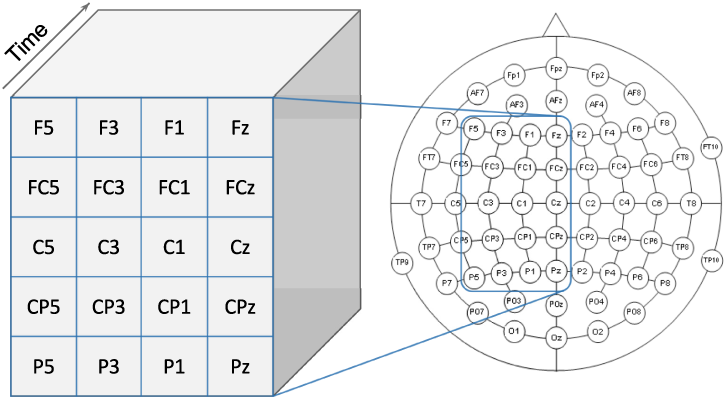
Input data arrangement for the ConvNet based on the spatial distribution of the recorded channels in healthy volunteers

#### 2.1.1. Pre-processing

EEG was notch-filtered (50 Hz) using a zero-phase filter in order to reduce power line interference and the baseline (1-s interval before the cue) was subtracted from all trials. No further pre-processing or filtering was applied to the EEG signals, and noisy epochs were not removed, in order to minimise user bias. Forty trials per task were executed, resulting in 160 trials per subject. Trials were subsequently segmented into 500-ms epochs, from 600 ms to 100 ms before movement onset (Fig. 2). EEG epochs were finally arranged in a 5 *×* 4 *×* 250 matrix with a two-dimensional spatial distribution and the time samples in the third dimension (Fig. 1). Data were divided in 128 trials (80%) for training and validation and 32 trials (20%) for testing. The training and validation set was further split into 102 trials (80%) for training and 18 trials (20%) for validation.

**Figure 2:**
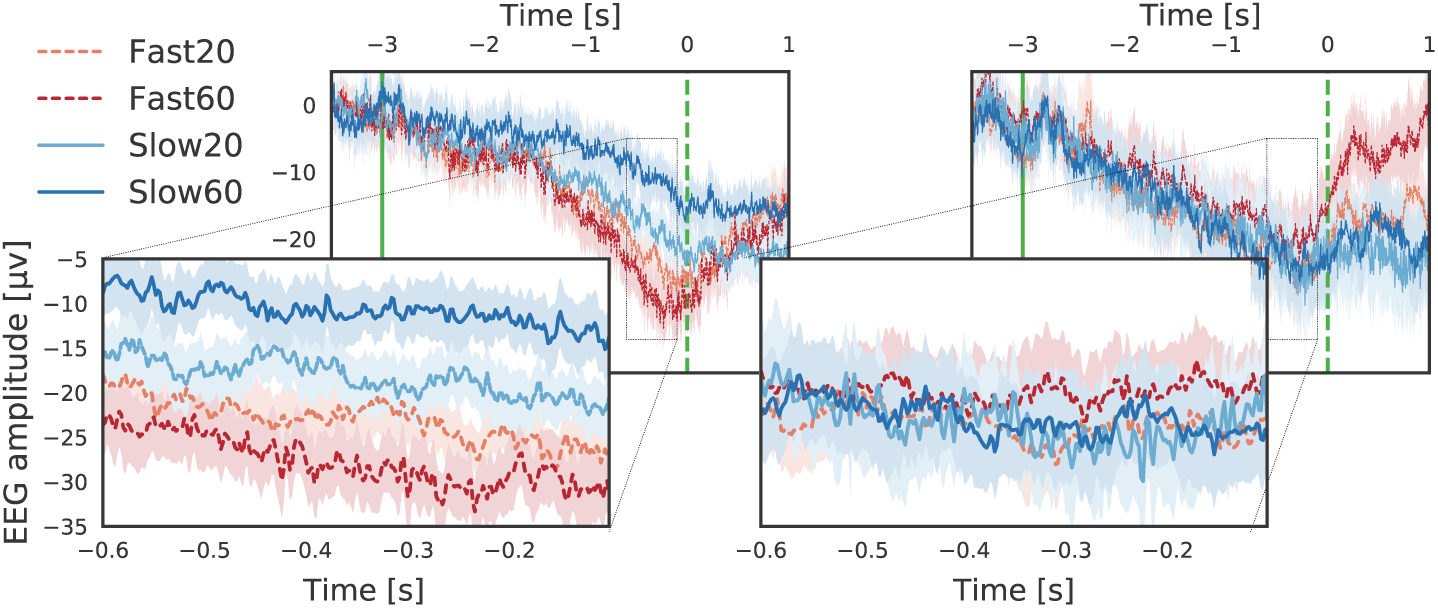
Representative examples of averages of 3 s EEG trials (back) and the corresponding 500 ms epochs (front) for healthy volunteers relative to movement onset. The solid trace and shading represent mean and 95% confidence intervals for each class, respectively, derived using 5000 bootstrap iterations. Vertical lines represent cue (solid) and movement onset (dashed) times.

### 2.2. Prediction strategies

#### 2.2.1. Convolutional Neural Network

The model was based on the EEGnet described in Lawhern et al. [21]. The ConvNet was built in TensorFlow 1.11 [22] using the Keras API [23] and trained on an Dell Precision 7910 workstation with an NVIDIA Titan Xp GPU, using CUDA 9 and cuDNN 7.3.

The model consisted of two blocks (Table 1). The input of the first layer was a pre-processed three-dimensional (3D) matrix for each trial, which was reshaped to apply four temporal filters (*F*_1_) to each channel. Following the original net architecture, convolutional kernels of size (1, 64) were applied in the temporal dimension. Kernel weights were initialized with a Glorot uniform technique, without applying a bias vector. The spatial dimension size was kept constant through zero padding without stride. Then, a batch normalization was applied. Afterwards, the matrix was reshaped and its dimensions were permuted in order to apply a depthwise convolution to every temporal slice by means of the wrapper time distributed layer [24]. Two spatial filters of size (*Cx, Cy*) for each feature map (depth multiplier parameter *D*) were applied and then the matrix was reshaped and dimensions were permuted again. Afterwards, a batch normalization followed by a Exponential Linear Unit (ELU) activation with alpha = 1, an average pooling of size (1, 4), and drop-out with a rate of 0.25 were applied.

**Table 1:** ConvNet architecture.

| Block | Layer | Output size | Param. # |
| --- | --- | --- | --- |
| 1 | Input | $(C_y \times C_x \times T)$ | 0 |
| | Reshape | $(1 \times C_y * C_x \times T)$ | 0 |
| | Conv2D $(1, 64) \times F_1$ | $(4 \times C_y * C_x \times T)$ | 256 |
| | Batch normalization | $(4 \times C_y * C_x \times T)$ | 16 |
| | Reshape | $(4 \times C_y \times C_x \times T)$ | 0 |
| | Permute | $(T \times 4 \times C_y \times C_x)$ | 0 |
| | TD $(C_y, C_x) \times D * F_1$ | $(T \times D * F_1 \times 1 \times 1)$ | 160 |
| | Permute | $(D * F_1 \times 1 \times 1 \times T)$ | 0 |
| | Reshape | $(D * F_1 \times 1 \times T)$ | 0 |
| | Batch normalization | $(D * F_1 \times 1 \times T)$ | 32 |
| | Activation (ELU) | $(D * F_1 \times 1 \times T)$ | 0 |
| | AveragePooling2D (1, 4) | $(D * F_1 \times 1 \times T/4)$ | 0 |
| | Dropout (.25) | $(D * F_1 \times 1 \times T/4)$ | 0 |
| 2 | SeparableConv2D (1, 16) $\times F_2$ | $(F_2 \times 1 \times T/4)$ | 192 |
| | Batch normalization | $(F_2 \times 1 \times T/4)$ | 32 |
| | Activation (ELU) | $(F_2 \times 1 \times T/4)$ | 0 |
| | AveragePooling2D (1, 8) | $(F_2 \times 1 \times T/32)$ | 0 |
| | Dropout (.25) | $(F_2 \times 1 \times T/32)$ | 0 |
| | Flatten | $(F_2 * T/32)$ | 0 |
| | Dense | $(N)$ | 228 |
| | Activation (Softmax) | $(N)$ | 0 |
| Total |  |  | 916 |

$C_x$ = channels (mediolateral direction), $C_y$ = channels (anteroposterior direction),
$T$ = time samples, $F_1$ = number of temporal filters, TD = TimeDistributed (DepthwiseConv2D), $D$ = depth multiplier (number of spatial filters), $F_2$ = number of pointwise filters, $N$ = number of classes.

In the second block, a 2D separable convolution of size (1, 16) with eight filters (*F*_2_) was applied. Henceforth, ELU activation, batch normalization, average pooling of size (1, 8), and drop-out were applied using the same hyperparameters as in the first block. Finally, the data was flattened to a single dimension and the four resulting scores of the dense layer were transformed to probabilities by means of a softmax activation.

The learning process consisted of a fixed number of learning steps using mini-batchs of 16 randomly selected trials and the Adam optimization. The initial number of learning steps was set to 500, and validation accuracy and loss curves as a function of the number of learning steps were obtained in order to derive the smallest number of learning steps required to achieve an acceptable classification accuracy. The loss obtained from the validation set was used as metric, and the model was updated if the loss decreased compared to the last saved model. To prevent model overfitting, only the model with the lowest validation loss was kept. In this regard, the relationship between training set size and performance was also analyzed to verify that the training set size was appropriate in relation to the dataset size [25].

#### 2.2.2. Alternative ConvNet configurations

The original ConvNet architecture presented in [21] was devised taking into account temporal and spectral characteristics of the EEG signals recorded for that particular study, such as sampling rate, window size and frequency resolution of the filters resulting from the convolutional kernels. For this reason, an alternative architecture was also tested here, whose parameters were derived from extrapolating the original criteria to match the characteristics of the dataset used in this study. The resulting architecture had the first convolutional kernels of size (1, 250), an average pooling of size (1, 4) in the second block. The rest of the parameters remained unchanged. Furthermore, we also explored the effect of the EEG window onset and length: in addition to the EEG epoch configuration described above, we also tested four alternatives: 500-ms epochs, from 1600 to 1100 ms before movement onset; 500-ms epochs, from 1100 to 600 ms before movement onset; 1000-ms epochs, from 1100 to 100 ms before movement onset; and finally, 1500-ms epochs, from 1600 to 100 ms before movement onset.

#### 2.2.3. State-of-the-art prediction strategies

Three state-of-the-art prediction strategies were selected, in order to compare their performance against ConvNets: support vector machines (SVMs), and decision trees using the boostrap aggregating (TB) and random forest ensemble algorithm (RF).

##### Support vector machines

SVMs are popular supervised learning models used for classification, that transform a non-linearly separable problem into a linearly separable problem by projecting data into a new feature space through the use of kernel functions, in order to find the optimal decision hyperplane in this feature space. This method was initially proposed to solve two-class problems, although strategies were later developed to extend this technique to multi-class classification problems [26]. For this study, SVMs were implemented for reproducibility purposes, as they would allow a direct comparison with prior studies using the same data [15, 16]. With regards to the SVM parameters, a Radial Basis Function (RBF) were used as kernel. Based on a heuristic search, the cost hyperparameter of the SVM was set to 0.001 and the gamma hyperparameter of the kernel was set to 0.0002. Furthermore, a one-against-one strategy was used to implement the multi-class SVM prediction strategy. This method constructed *k(k-1)/2* classifiers, where *k* is the number of classes of the problem. Each classifier used the training data from two classes chosen out of *k* classes. After the training process was over, a voting strategy was used to determine to which class each pattern belongs to. The open source library tool for classification and regression problems (LIBSVM) was used to build the SVMs [27].

##### Decision trees

a decision tree is a non-parametric technique used in regression and classification problems, that makes sequential, hierarchical decision about the outcomes variable based on the predictor data. In particular, the bagging algorithm attempts to improve this classification by combining classifications of randomly generated training sets. Briefly, it consists of generating multiple versions of a predictor and using these to get an aggregated predictor [28]. Many bootstrap replicas of this dataset and decision trees on these replicas must be generated to bag a weak learner such as a decision tree on a dataset. Then, each bootstrap replica is obtained by randomly selecting N observations out of N with replacement (where N is the dataset size). Bagging works by training learners on resampled versions of the data. This resampling is usually done by bootstrapping observations, that is, selecting N out of N observations with replacement for every new learner. In addition, every tree in the ensemble can randomly select predictors for decision splits [29]. An average of predictions is taken from individual trees to find the predicted response of a trained ensemble. Aggregation averages the versions when predicting a numerical outcome and performs a plurality vote when predicting a class. The multiple versions are created by making bootstrap replicas of the learning set and using them as new learning sets. On the other hand, random forest is an ensemble-based learning technique that trains many weak learners (such as decision trees) to solve the algorithm with bootstrap aggregation technique. This method uses random feature selection in the tree induction process and finally aggregates the predictions of the ensemble to make the final decision. When a new object is to be classified from an input vector, it passes the sample vector to each of the trees in the forest so that each tree can provide a classification decision, and then results of individual trees are combined to choose the classification [30].

Unlike ConvNets, these methods require a separate feature extraction stage before classification [31, 32, 16, 33]. In this regard, ten features were calculated for each 500-ms epoch: 1) Basal amplitude value, using the Hilbert transform to estimate the area envelope, 2) Kurtosis, 3) Curve length, as the sum of consecutive distances between amplitudes, 4) Noise level, as 3 times the standard deviation of the amplitudes, 5) Number of positive peaks, 6) Average nonlinear energy, using the Teager energy operator, 7) Number of zero crossings, 8) Maximum negativity peak, 9) Root-mean-square (RMS) amplitude, and 10) Average power in the interval from 0 to 5 Hz, using Welch power spectral density estimator with a Hamming window and a 50% overlap.

### 2.3. Data analysis

An individual prediction strategy was trained for each subject, and the same data partitioning for training, validation, and testing was used for all prediction strategies. Additionally, ConvNets were trained with the same dataset partitioning, but with randomly scrambled labels, in order to determine the chance classification accuracy level. Furthermore, ConvNets were also trained with increasing number of examples in order to test the effect of training set size on classification accuracy. To ensure the generalizability of the final results, a 5-fold cross-validation procedure was performed, in which a new individual prediction strategy was trained for each fold. The overall classification accuracy and Cohen’s *κ* (a metric that compares observed accuracy with expected accuracy due to random chance), together with per-class precision and recall were quantified for each subject, as the mean values of the 5-fold cross-validation procedure computed from the test sets. Finally, to ensure the reproducibility of these results, the source code for the ConvNet is available at https://github.com/ragatti/STSnet and the dataset is available from the corresponding author upon request.

### 2.4. Statistics

The Shapiro-Wilk test was performed in order to assess the assumption of normality, which held for all indexes. Performance indexes are reported as *mean ± standard deviation* unless stated otherwise. A paired t-test was used to assess differences in overall classification accuracy and Cohen’s *κ* between ConvNet architectures, and to evaluate differences between different epoch onsets and lengths. A repeated measures analysis of variance (RMANOVA) was used to assess differences in accuracy, *κ*, precision and recall, with *Classifier, Architecture, Speed* and *Force* as factors (whenever applicable). Main effects and two-way interactions were analysed, and the Greenhouse-Geisser correction was applied to account for deviations in sphericity. Furthermore, the Tukey test was used for post hoc comparisons. In line with current statistical trends [34], no fixed threshold for statistical significance was set, and the results are otherwise analysed in terms of the effect sizes and their experimental relevance.

## 3. Results

### 3.1. ConvNet validation

The evolution of the validation accuracy and validation loss as a function of the number of learning steps is shown in Fig. 3 (left). It can be observed that the accuracy reaches a stable value after 100 steps while the loss stabilises after approximately 300 steps, indicating that more training steps would not improve the results and that the ConvNet is not overfitting the data. As a reference, training the ConvNets using 500 steps took approximately 3 min per subject and classifying each new trial took approximately 7 ms. Furthermore, Fig. 3 (right) shows the chance level accuracy obtained after training the model with randomly scrambled labels. In this case, the prediction accuracy is close to the theoretical chance level of 25% for a 4-class classification problem and it does not improve with the number of training steps.

**Figure 3:**
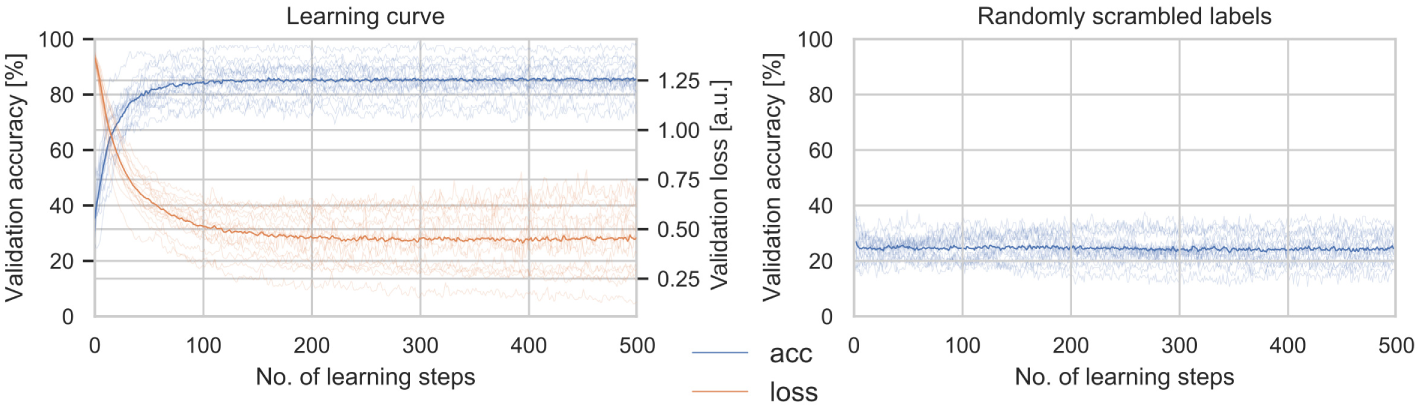
Left: evolution of the validation accuracy (acc) and validation loss (loss) from all volunteers as a function of the number of learning steps. Right: Validation accuracy from all volunteers as a function of the number of learning steps with randomly scrambled labels. In both the dark line represents the mean validation accuracy for all subjects in each group, and each light line represents the mean validation accuracy for a single subject, derived from the 5-fold cross-validation.

Fig. 4 shows the relationship between test accuracy and training set size, from which it can be deduced that the ConvNet strategy can be trained with as few as 80 examples and still achieve an acceptable classification accuracy above 80%.

**Figure 4:**
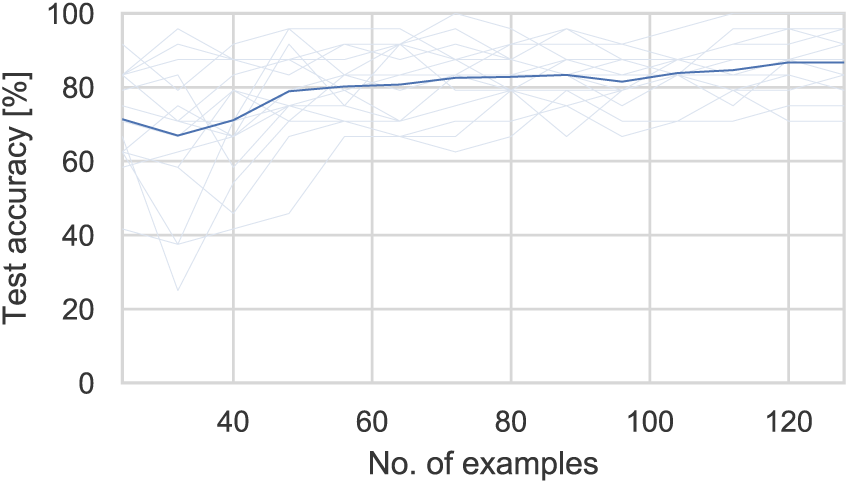
Test accuracy from all healthy volunteers as a function of the number of examples. The dark line represents the mean test accuracy for all subjects in each group, and each light line represents the mean test accuracy for a single subject, derived from the 5-fold cross-validation.

### 3.2. Alternative ConvNet configurations

No differences in accuracy and *κ* were found between the original (84.0±7.0% and 0.79 ± 0.09) and the alternative architecture (83.8 ± 5.6%; *t*_15_ = 0.200, *p* = 0.844 and 0.78 ± 0.07; *t*_15_ = 1.206, *p* = 0.246). Considering that the alternative architecture required over 500 additional parameters compared to the original, we decided to use the latter for all further comparisons. Furthermore, Table 2 shows the performance of the selected architecture, in which EEG epochs with different onset and lengths were compared against the default epoch configuration (500 ms duration, from 0.6 to 0.1 s before movement onset). We did not observe significant improvements due to window onset or length in comparison to the proposed epoch configuration; however, it is worth noting that the window from 1.6 to 1.1 s before movement onset showed a consistently worse performance, which might indicate that there is additional discriminative information for classification located in the interval between 1.1 s and movement onset.

**Table 2:**
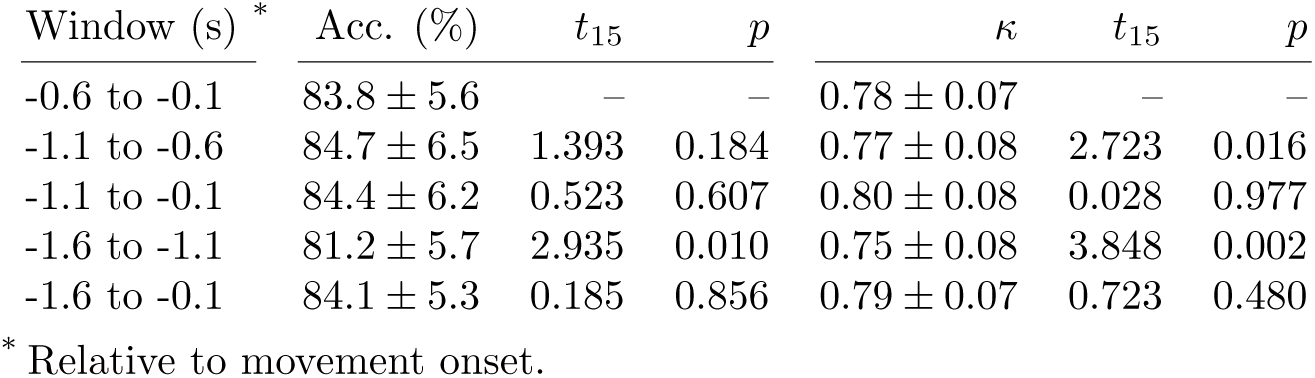
Performance of the ConvNet architecture as a function of window onset and length.

| Window (s) * | Acc. (%) | $t_{15}$ | $p$ | $\kappa$ | $t_{15}$ | $p$ |
| --- | --- | --- | --- | --- | --- | --- |
| -0.6 to -0.1 | $83.8 \pm 5.6$ | — | — | $0.78 \pm 0.07$ | — | — |
| -1.1 to -0.6 | $84.7 \pm 6.5$ | 1.393 | 0.184 | $0.77 \pm 0.08$ | 2.723 | 0.016 |
| -1.1 to -0.1 | $84.4 \pm 6.2$ | 0.523 | 0.607 | $0.80 \pm 0.08$ | 0.028 | 0.977 |
| -1.6 to -1.1 | $81.2 \pm 5.7$ | 2.935 | 0.010 | $0.75 \pm 0.08$ | 3.848 | 0.002 |
| -1.6 to -0.1 | $84.1 \pm 5.3$ | 0.185 | 0.856 | $0.79 \pm 0.07$ | 0.723 | 0.480 |

\* Relative to movement onset.

### 3.3. Performance of the classification strategies

Overall performance in terms of accuracy and *κ* was affected by the choice of *Classifier*, as shown in Fig. 5 (One-way RMANOVA; *F*_3,45_ = 27.35, *p* < 0.001 and *F*_3,45_ = 29.63, *p* < 0.001, respectively). Specifically, ConvNets showed higher accuracy and *κ* values compared to all other strategies (Tukey; *p* < 0.001 for both indexes), which in time did not show any relevant differences among them (Tukey, *p* > 0.985). With regards to per-class classification indexes (Fig. S1), precision and recall were likewise only affected by the choice of *Classifier* (Three-way RMANOVA; *ϵ* = 0.53, *F*_1.6,24.1_ = 28.77, *p* < 0.001 and *ϵ* = 0.55, *F*_1.6,24.7_ = 26.58, *p* < 0.001, respectively). No other main effect or interactions significantly affected per-class indexes. Post hoc analysis showed that ConvNets outperformed all other strategies (Tukey, *p* < 0.001 for both indexes), which did not show any relevant differences among them either (Tukey, *p* > 0.894).

**Figure 5:**
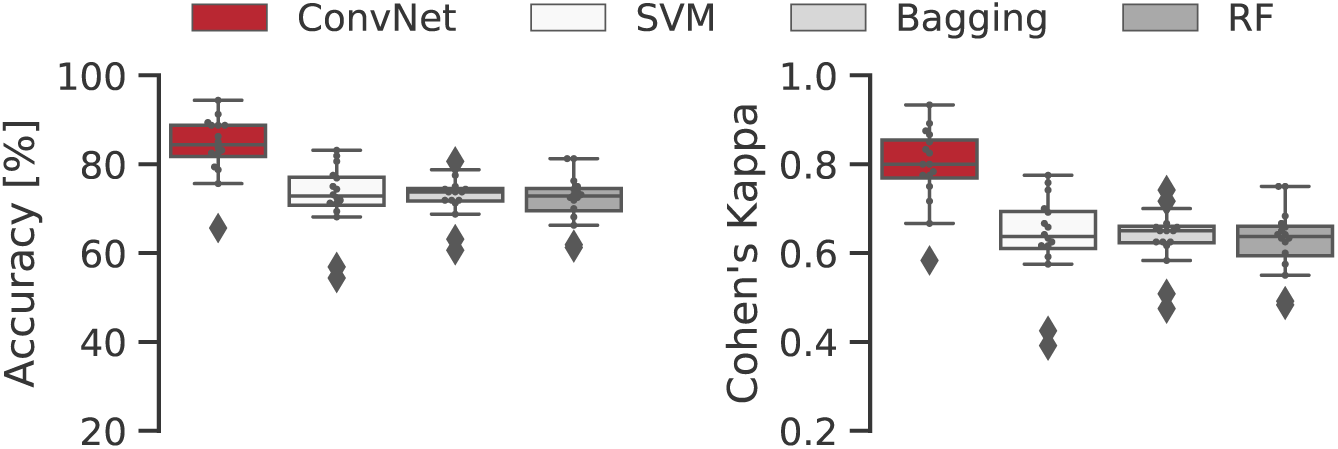
Classification results over test data for different strategies. Accuracy and *κ* for healthy volunteers (n = 16). Boxes represent the median and the 25^th^ and 75^th^ percentiles, whiskers represent 5^th^ and 95^th^ percentiles, diamonds represent values outside of the 5^th^ - 95^th^ percentile range and the individual dots represent the average Accuracy/*κ* for each individual subject, calculated from the 5-fold cross-validation.

### 3.4. ConvNet architecture analysis

Fig. 6 shows the resulting network architecture for a randomly selected subject (S4). We attempted to find common patterns across subjects in the temporal and spatial filter layers, but we did not observe any specific features that may relate to specific neurophysiological patterns, and we did not detect any patterns that could be generalised across subjects either. However, we did identify a common pattern across subjects in the 2D separable convolution layer. Indeed, Fig. 7 shows that the kernels in this layer have weights that are either predominantly positive (in blue) or negative (in red) along the temporal direction (represented as columns in Fig. 7). We interpreted this as a sign that the spatial information is more relevant for classification than temporal information. We also observed that the kernels were very similar in each layer, which might be signifying a certain degree of redundancy in terms of the number of filters and depth.

**Figure 6:**
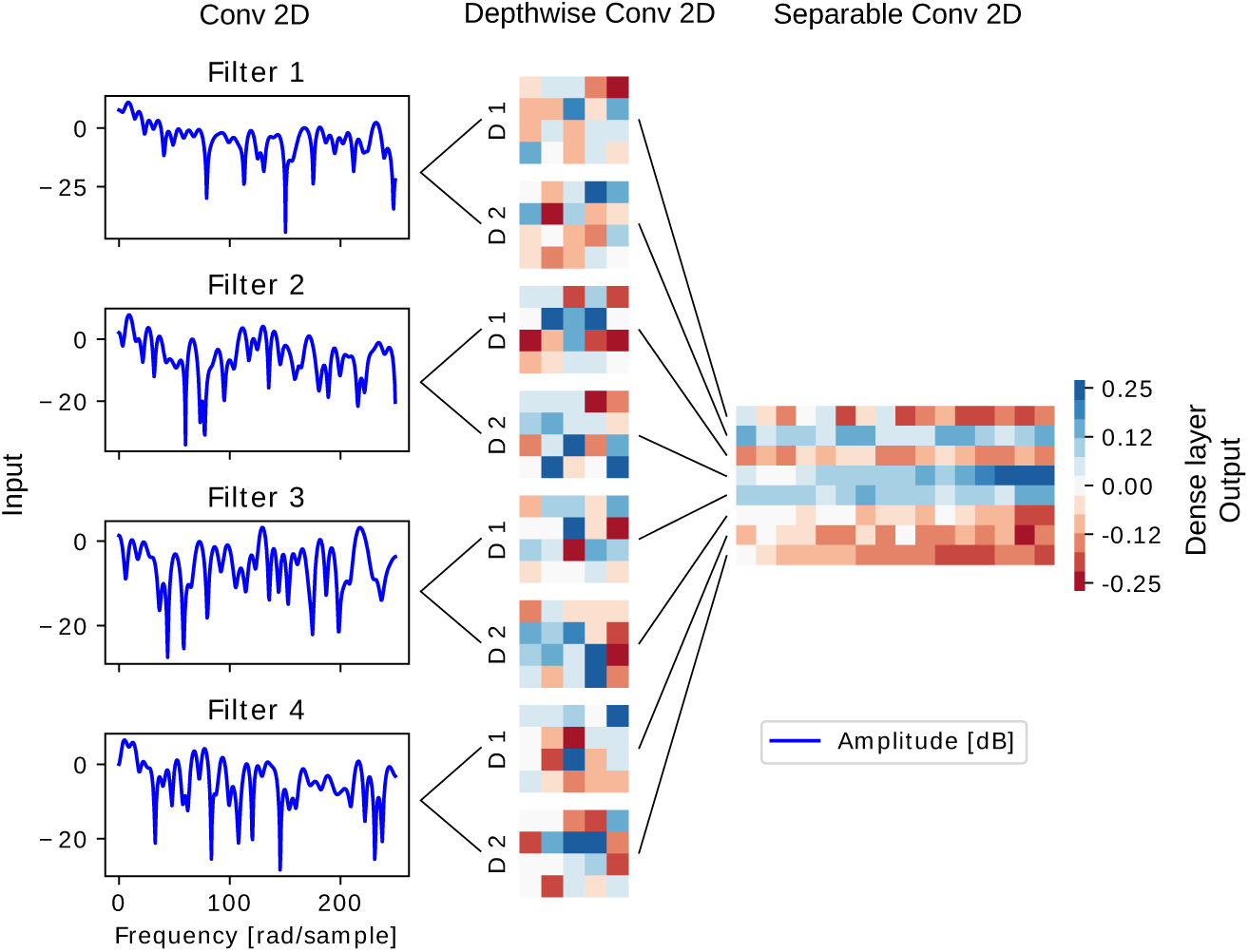
Example of resulting ConvNet architecture for a randomly selected subject (S4). Left: frequency response of the four temporal filters of the first layer. Center: spatial kernels for the intermediate layer. Right: 2D separable convolution kernel.

**Figure 7:**
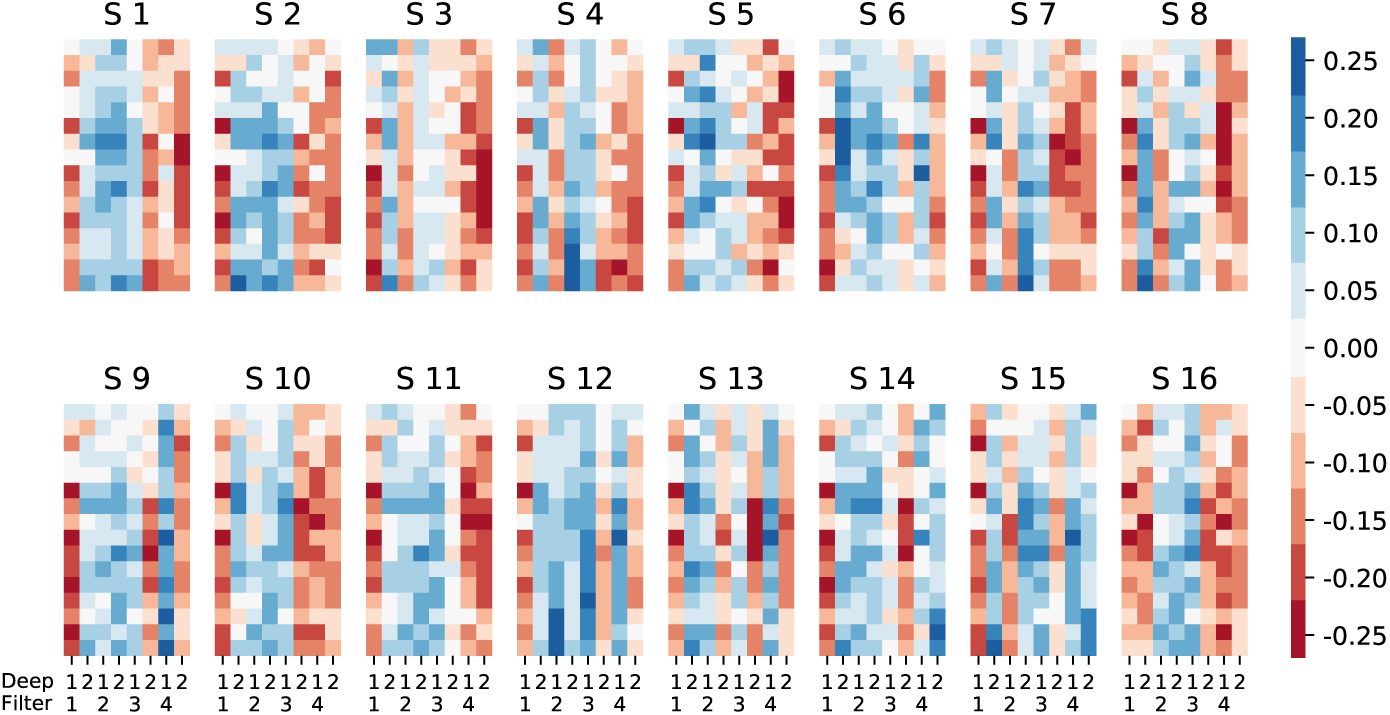
Kernels from 2D separable convolution layer for all volunteers. Each ConvNet has 8 spatial kernels (columns) with 16 temporal values each (rows) in this layer.

In light of this, we tested the performance of three new architectures: 4 temporal filters with a depth multiplier of 1 (ConvNet-4,1), 2 temporal filters with a depth multiplier of 2 (ConvNet-2,2) and 2 temporal filters with a depth multiplier of 1 (ConvNet-2,1), against the original configuration (ConvNet-4,2). These architectures presented different numbers of parameters: ConvNet-4,2 had 916 parameters in total, from which 876 were trainable, whereas ConvNet-4,1 had 580 parameters (556 trainable), ConvNet-2,2 had 444 parameters (424 trainable), and finally, ConvNet-2,1 had 288 parameters (276 trainable).

We observed that accuracy and *κ* depended on the selection of *Architecture*, as shown in Fig. 8 (One-way RMANOVA; *F*_3,45_ = 28.17, *p* < 0.001 and *F*_3,45_ = 38.76, *p* < 0.001, respectively). The post hoc analysis revealed that there were no differences in performance between ConvNet-4,2, ConvNet-4,1 and ConvNet-2,2 (Tukey, *p >* 0.971), and that these three strategies out-performed ConvNet-2,1 (Tukey, *p* < 0.001). Furthermore, per-class classification indexes (Fig. S2), showed the same behaviour. Essentially, *Architecture* was the only significant factor for precision (Three-way RMANOVA; *ϵ* = 0.67, *F*_2.1,30.2_ = 29.08, *p* < 0.001) and recall (Three-way RMANOVA; *ϵ* = 0.64, *F*_1.9,28.8_ = 28.18, *p* < 0.001). No other main effect or interactions significantly affected per-class indexes. Post hoc analysis showed that ConvNet-2,1 performed worse than all other strategies (Tukey, *p* < 0.001 for both indexes), with no significant differences among them (Tukey, *p* > 0.971).

**Figure 8:**
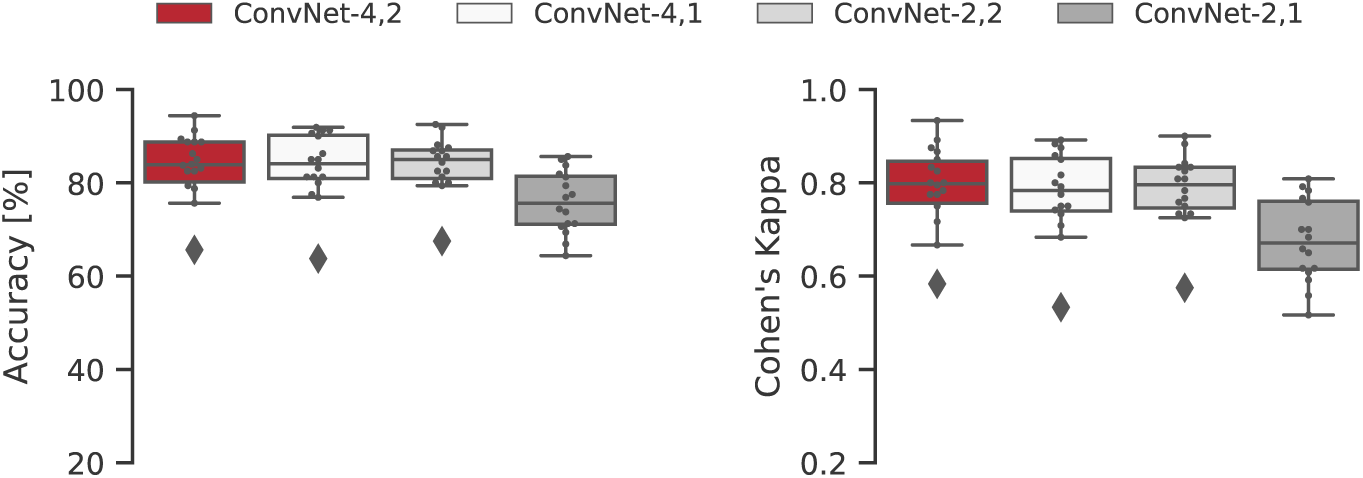
Classification results over test data for different architectures. Accuracy and *κ* for healthy volunteers (n = 16). Boxes represent the median and the 25^th^ and 75^th^ percentiles, whiskers represent 5^th^ and 95^th^ percentiles, diamonds represent values outside of the 5^th^ - 95^th^ percentile range and the individual dots represent the average Accuracy/*κ* for each individual subject, calculated from the 5-fold cross-validation.

## 4. Discussion

### 4.1. Neurophysiological aspects of movement prediction

Building efficient movement decoding models from brain signals is crucial for many biomedical applications, particularly in the BCI field that require precision in online control of assistive devices. Moreover, decoding specific movement features, such as speed, force and/or direction, provides additional degrees of freedom, resulting in more accurate and natural motor commands at the expense of increasing the complexity of the decoding problem [35, 14, 12, 13]. Early attempts to decode movement from brain signals during movement execution or imagination were focused on classifying between limb movements [36, 37, 38, 39]. Classification accuracy for these studies was close to 80% for 2 classes [36, 37], and close to 75% for 4 classes [39]. Other studies have tried to decode movement of specific body parts from a single limb, such as wrist [40], or individual finger movements [9], obtaining similar results.

On the other hand, prediction of movement, i.e., decoding movement not during, but before its execution, is a much more difficult task. Considering the brain as a predictive neural system, expectation can be seen as a representation of prediction that serve to sensory or motor areas as preparatory processing prior to an event, particularly in short time scales [41]. Movement intention is the first interesting command to decode from EEG before a movement is executed, as trigger for other more complex motor instructions. It is well known that information about movement intention is encoded in the MRCPs, around 1.5 s prior to movement onset [7]. The timing of the prediction is a relevant feature to study, since it has been shown that a sensory stimulus delivered synchronously with the peak negativity of the MRCP maximises neural plasticity [42]. Furthermore, kinetic information encoded in the movement intention could be particularly useful; for example, by decoding these movement parameters it would be possible to introduce task variability in the rehabilitation training, which has been shown to maximise the motor learning [43].

It has been already shown that pre-movement EEG contains valuable information about motion. Indeed, detection of voluntary movement from single trial EEG using a matched filter approach demonstrated relatively good performance in a 2-class classification scheme (sensitivity *≈* 82.5% for healthy subjects) [44]. However, classification rates for multi-class classification problems are still relatively low in healthy volunteers. As an example, recent studies directed towards the extraction of additional information from movement intention beyond simple detection, such as the prediction of the body part that is about to perform the movement [18], or the classification between different types of movement used in daily life, such as palmar, lateral and pinch grasps [10], resulted in classification accuracies not better than chance levels for the 4-class classification attempts.

In particular, previous work with the same dataset used in this study obtained mean accuracy values of approximately 32-40% for the 4-class classification [16], which is on par with the chance level for that type of problem [45]. These result might be partially explained by the fact that the aim of the study was to obtain a fast prediction scheme using few electrodes and a simple classifier that did not require extensive calibration. As such, only one channel was used as input, and the signals were band filtered using low cut-off frequencies values. However, it was recently suggested that information from the entire EEG spectrum is needed to discriminate between task-related parameters from single-trial movement intention [17].

Based on this idea, in this study it was possible to significantly improve the movement prediction accuracy using twenty available channels without additional pre-processing, such as artifact removal or epoch selection. Accuracy levels reached values close to 85% in healthy volunteers, representing an improvement of almost 45% compared to previous results. Furthermore, previous studies have shown that classification of speed tasks achieved higher accuracy in the prediction of ankle dorsiflexion movements [15]. However, it was not the case for hand grasping tasks, since no significant effects of speed or force and no interactions were observed, in line with previous results [16].

In light of this, it could be hypothesised that the decoding of complex movement requires more information (in terms of number of channels or features) in order to achieve a classification accuracy comparable with that obtained for simpler movements, such ankle or wrist flexion/extension (binary classification problems) [46, 40, 47, 15]. In this regard, we observed that neither shifting nor increasing the length of the time window improved the ConvNet performance (Table 2). We also altered the original ConvNet architecture to better match the time-frequency characteristics of our database, again without a significant improvement in performance (Fig. 8). Finally, the analysis of the ConvNet architecture suggested that the network performs a complex integration between time-frequency and spatial features of the EEG signal, and that the high channel density over a relatively small scalp area in this particular dataset might be crucial in order to reach a high classification accuracy.

### 4.2. Methodological aspects of movement prediction

Deep learning methods were originally developed in the computer vision field [48], and recently gained popularity in EEG analysis, in which they are used with the aim of improving classification performance over more traditional approaches, such as linear discriminant analysis, k-nearest neighbours or SVMs [20]. ConvNets are a type of feed-forward deep learning networks that are useful when data have a known topological structure [19, 25]. As a representation learning method, one of the advantages of ConvNets is that feature extraction and classification is intrinsically optimised. Typically, ConvNets consist of a combination of convolutional and pooling layers. The convolutional layer applies mathematical convolution operations through a number of kernels that perform a local weighted sum along the input and return each one in a feature map. Then, the same weights are shared across the input and have only local connections, thereby reducing the amount of network parameters. The pooling layer performs a reduction of the input by applying a function to nearby units, e.g. the maximum value among neighbours, where the units are the pixels of an image or the temporal samples of a biosignal.

The ConvNet implemented in this study is based on a recently proposed architecture that demonstrated good performance employing a small number of parameters in the classification of EEG signals recorded using different paradigms [21]. In this model, the first convolutional layer works as a frequential filter, in which the outcome consists of four different band-pass filters that minimise the error at the output. In accordance with the input structures used in image processing, the EEG input to a ConvNet is usually reshaped into a 2D distribution, by arranging channels along the rows and time samples in the columns [49, 50] or by transforming the input into a new space [8], e.g., to a time-frequency domain through Fourier transform and averaging along the channels [51, 52]. The ConvNet implemented in this study considered the localisation of the electrodes in order to keep the spatial relationship between them. Furthermore, EEG signals are commonly pre-processed by using temporal and spatial filters, and epochs containing artifacts or with amplitudes above a certain threshold are rejected in order to improve the signal-to-noise ratio [7]. These processes are time-consuming, prone to user bias and may result in the loss of useful information to decode movement. Taking this into consideration, only minimal and automatic pre-processing (baseline correction and notch filtering) was performed in this study prior to the classification stage, and no epochs were removed. Furthermore, it is worth noting that this strategy does not require extremely large datasets for training and the training time is negligible compared to the average setup time for a BCI, which makes it viable for use in rehabilitation.

Since the main goal of this work was to determine the achievable levels of prediction accuracy from single-trial EEG using state-of-the-art machine learning techniques, we compared the results obtained using ConvNets with state-of-the-art prediction strategies. The first comparison is with a strategy based on SVMs, to allow for a direct comparison with previously published results [15, 16]. The prediction results of the ConvNet were better than the SVM for all tasks and all performance indexes in healthy volunteers by an average of 12 percentage points (Fig. S2). This is even more relevant considering that the SVMs implemented in this study (using twenty available channels) already improved the classification accuracy by approximately 32 percentage points compared to the previous study with the same dataset (using only a single channel, C3, plus an eight-channel Laplacian filter) [16]. Furthermore, the performance of classification strategies based on decision trees was also significantly lower compared to ConvNets. Indeed, accuracy and *κ* values obtained with decisions trees were not different from those obtained with SVMs, using the same features.

Furthermore, a systematic investigation regarding movement prediction performed with combinations of spatial filtering (principal component analysis, independent component analysis, common spatial patterns analysis, and surface Laplacian derivation), temporal filtering (power spectral density estimation and discrete wavelet transform), pattern classification (linear and quadratic Mahalanobis distance classifier, Bayesian classifier, multi-layer perceptron neural network, probabilistic neural network, and SVM), and multivariate feature selection strategy using a genetic algorithm, achieved a maximum accuracy of 75% for binary classification [53]. Taken together, these results might imply that the differences in performance between ConvNets and other strategies are probably due to to the feature selection and generation methods.

In this regard, most of the current methods for feature selection and generation are defined by a human investigator based on *a priori* knowledge of the neurophysiology of the brain, for example in terms of time-frequency characteristics of the signals or how the sources of electrical activity are spatially distributed in the cortex. In fact, even standard EEG pre-processing (e.g. bandpass filtering or channel selection based on predefined brain activation patterns) could be inadvertently discarding relevant information for classification. On the other hand, ConvNets require minimal pre-processing and are not limited to feature selection or generation constrains. We changed the number of input features (by changing the window length), and tested several different architectures (by changing the number an parameters of the temporal and spatial filters), and the ConvNet still maintained excellent performance, even with a reduced number of parameters (83.9 ± 5.9% accuracy using 444 total parameters in ConvNet-2,2).

### 4.3. Limitations and future work

Several constraints need to be considered: attempts to use a single ConvNet to predict movements from all subjects resulted in low performance indexes during pilot tests (average accuracy of 27.4 ± 4.4%). This is not an issue in most real-life applications where the decoding is used to control a device for a single subject (and thus an individual ConvNet is trained for each subject), but nevertheless highlights the difficulty in describing a general behaviour of the EEG signal in terms of decoding force and speed. The same issue can be observed when attempting to understand and visualise of the specific features that allow a good classification, since it is not straightforward to extract and interpret physiological information from the network, and these feature vary between subjects. Furthermore, even if high accuracy was achieved offline, it is crucial to perform real-time tests with adequate feedback. Future work will be directed towards testing the strategy with a real application, for which an accurate detection of the movement onset is necessary and an idle state should be considered [54]. Finally, once the definitive scheme has been defined, efficient hardware implementations should be tested in chips or field-programmable gate arrays [19].

## 5. Conclusion

The results from this study suggest that hand movement speed and force can be accurately predicted from single-trial EEG using convolutional neural networks, although additional considerations are still required to transfer these protocols from laboratory to clinic.

## Acknowledgment

The workstation used in this study was provided by the Center for Neuroplasticity and Pain (CNAP), which is supported by the Danish National Research Foundation (DNRF121). The Titan Xp GPU used for this research was donated by the NVIDIA Corporation.

## Supplementary figures

**Figure S1:**
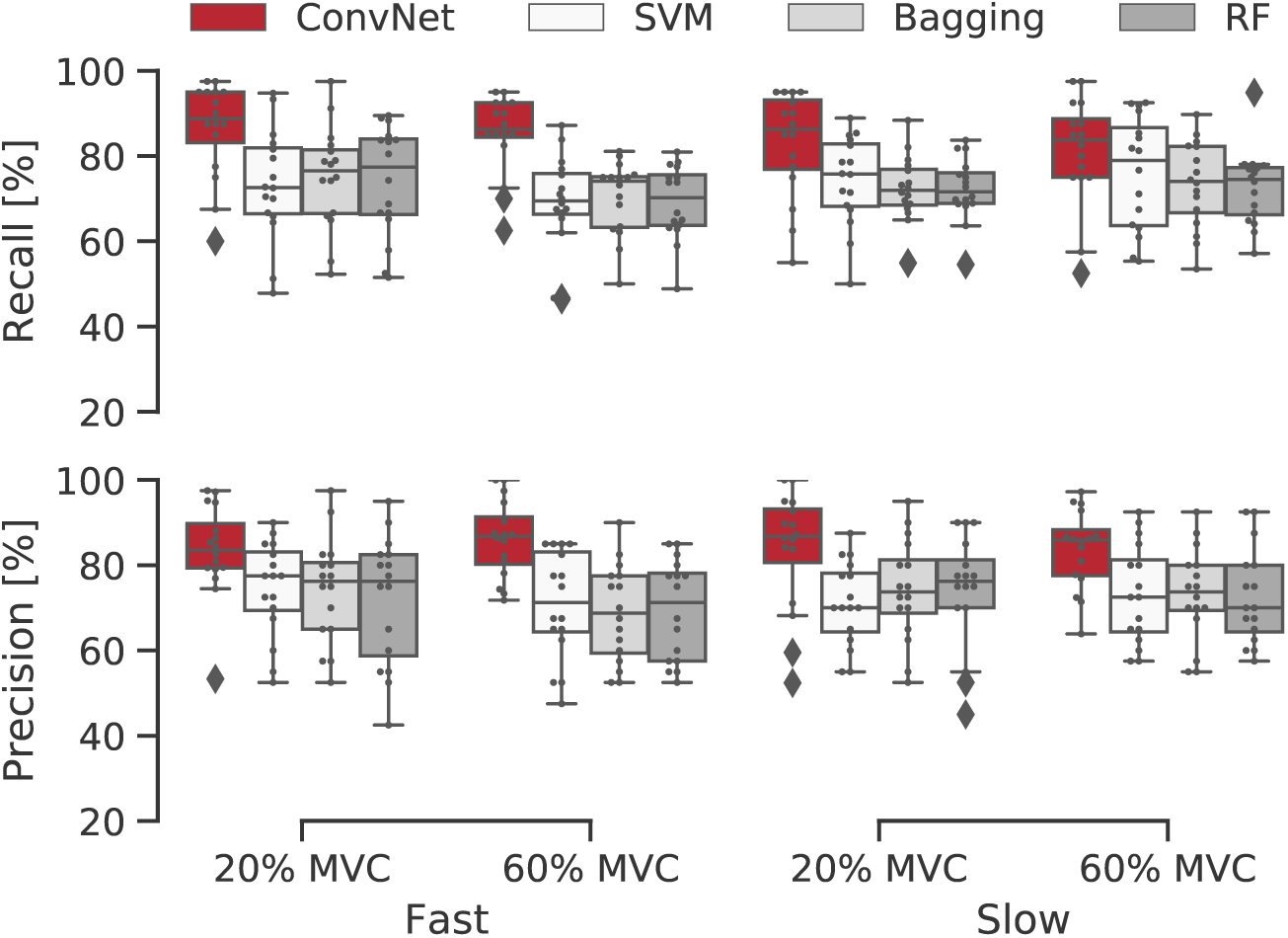
Classification results over test data for different strategies. Precision and recall for healthy volunteers (n = 16). Boxes represent the median and the 25^th^ and 75^th^ percentiles, whiskers represent 5^th^ and 95^th^ percentiles, diamonds represent values outside of the 5^th^ - 95^th^ percentile range and the individual dots represent the average precision/recall for each individual subject, calculated from the 5-fold cross-validation.

**Figure S2:**
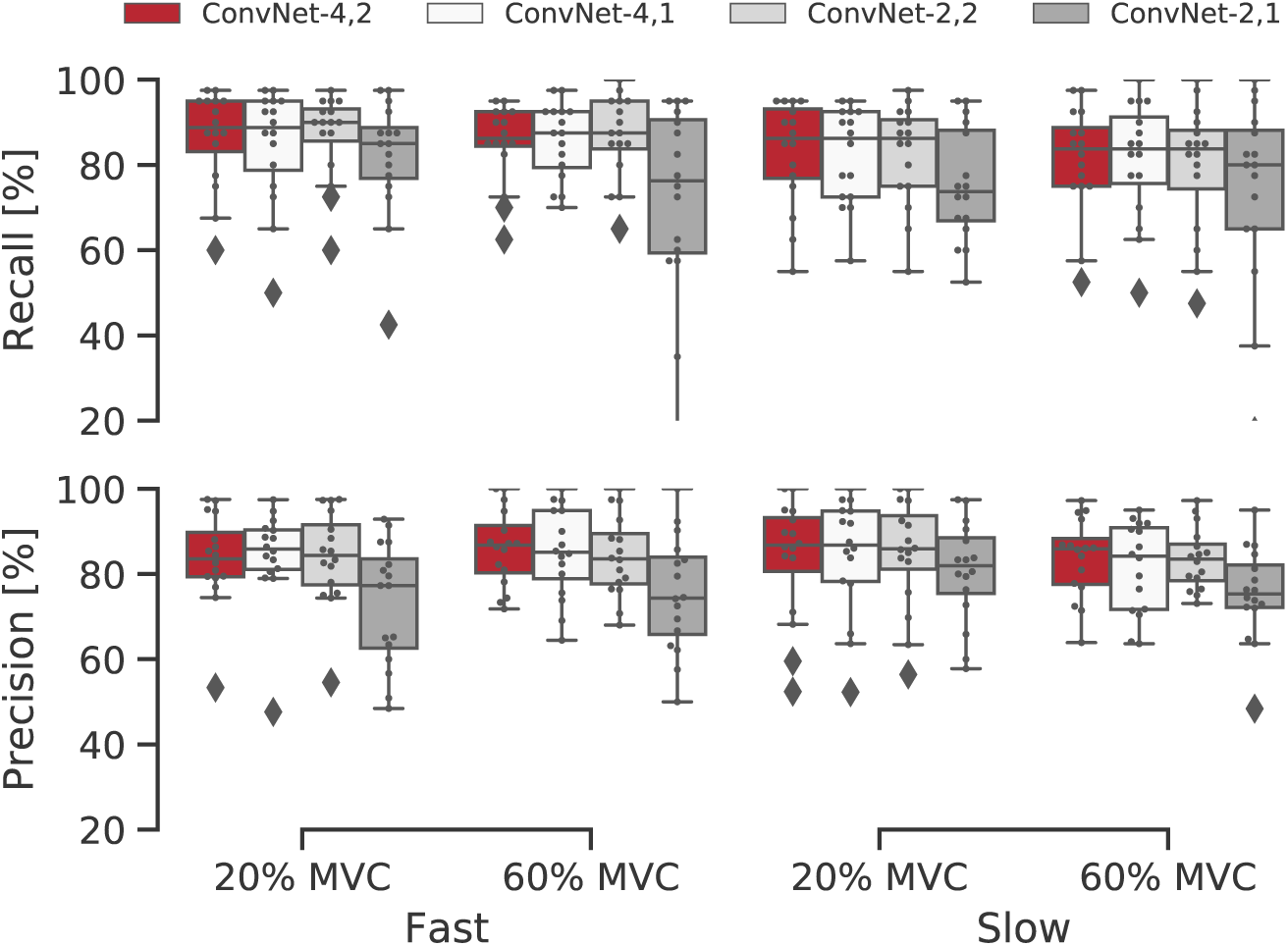
Classification results over test data for different architectures. Precision and recall for healthy volunteers (n = 16). Boxes represent the median and the 25^th^ and 75^th^ percentiles, whiskers represent 5^th^ and 95^th^ percentiles, diamonds represent values outside of the 5^th^ - 95^th^ percentile range and the individual dots represent the average precision/recall for each individual subject, calculated from the 5-fold cross-validation.

